# Real-time posterior probabilistic TMS motor mapping with robust individualized stopping criteria

**DOI:** 10.64898/2026.10.06.757035

**Authors:** Renxiang Qiu, Zi-Jian Feng, Ole Numssen, Yu-Feng Zang, Florian Wilhelmy, Erdem Güresir, Thomas R. Knösche, Konstantin Weise

## Abstract

**Background.** Electric-field-informed regression provides high-resolution transcranial mag-netic stimulation (TMS) motor maps, but conventional nonlinear fitting is too slow to update the map during acquisition, so sessions rely on predefined stimulation budgets, despite sub-stantial inter-individual variability in the data required for stable localization. **Objective.** To develop a real-time analysis updating a posterior probability map between pulses and compare its hotspots with finger-tapping functional MRI (fMRI). **Methods.** We analyzed two independent cohorts (12 and 13 participants, one newly collected); each hand muscle was analyzed independently. We replaced the conventional per-element sigmoid fit with Spear-man’s rank correlation coefficient and transformed the resulting score map into a posterior probability map using Bayesian evidence. Acquisition stopped once posterior mass within 5 mm of the hotspot passed a preset confidence level. **Results.** The Spearman and linear scores updated the region of interest within milliseconds, versus tens of seconds for sigmoid fitting. Spearman more closely matched the full-data sigmoid hotspot in recorded sessions and converged faster and more precisely to the true source in virtual subjects. Across both fast scores, at a confidence level of 0.85 the stopping criterion was met after a median of 74-159 pulses. FDI hotspots lay 8.3 ± 4.3 mm from the fMRI center of gravity, in the precentral hand region. **Conclusions.** Rank correlation reduces computation time without loss of accuracy relative to sigmoid fitting. The resulting posterior probability map provides a session-specific stopping criterion in place of a fixed pulse budget.

## 1. Introduction

Accurate localization of individual cortical muscle representations is important for studying human motor organization and for defining stimulation targets in experimental and clinical transcranial magnetic stimulation (TMS). Conventional motor maps identify the coil placements that evoke the largest motor evoked potentials (MEPs), but a coil placement does not uniquely specify the cortical tissue contributing to the response because each pulse induces a spatially distributed electric field. Electric-field-informed regression addresses this limitation by relating, for each stimulation, the field magnitude at every element of an indi-vidual cortical mesh to the observed MEP amplitude (Numssen et al., 2021; Weise, Numssen, et al., 2023; Weise et al., 2020). The field–MEP relationship is modeled as a linear function or a four-parameter sigmoid (Numssen et al., 2021; Weise, Numssen, et al., 2023) capturing the saturating recruitment curve (Devanne et al., 1997). The best-fitting element provides the estimated cortical representation.

Recent work has made this mapping more efficient through pre-experimental acquisi-tion design. The initial random-sampling study acquired 900–1100 pulses per participant and estimated retrospectively that approximately 180 pulses were required for a stable map (Numssen et al., 2021). The subsequent protocol recommends at least 300 (Weise, Numssen, et al., 2023). Prospective designs then chose the coil configurations before the session, either by sampling the space of electric-field patterns or by optimizing the whole pre-experimental sequence for field redundancy, cortical separability and coil travel (Qiu et al., 2026; Schultheiss et al., 2025). These approaches improve the choice of coil placements for the acquired pulses, but they do not determine from the accumulating data when an individual session has become sufficiently localized. The endpoint remains prescribed in advance or assessed retrospectively.

An efficient stopping criterion requires an interpretable measure of localization uncertainty. The coefficient of determination (*R*^2^) commonly used in the established map is a goodness-of-fit measure and does not quantify the evidence accumulated for a location. For example, the same *R*^2^ = 0.7 can arise from ten or one thousand pulses, although the two estimates do not carry the same statistical support. Ideally, this uncertainty would be ex-pressed as the probability that the true representation lies within a given cortical region, based on the evidence collected so far. This would allow us, for example, to define a criterion that requires a confidence of 85% in an area of 5 cm^2^ around the peak. Assessing convergence is also useful, because a predefined confidence limit may not be achievable, or because we may want to reach the highest achievable confidence level. Map stability and the stability of its peak were both proposed as online stopping criteria (Numssen et al., 2021). Both track only the change between successive estimates and can therefore stop early on a locally stable but incorrect hotspot (Section 3.3). These quantities must also be computed fast enough to update between pulses.

Recently, two Bayesian formulations have addressed the uncertainty of the mapping itself. Laine and colleagues update a spatial probability distribution after every stimulus under closed-loop multi-locus TMS, recording each response as either detected or not at a fixed amplitude threshold (Laine et al., 2026). They propose a stopping criterion based on the convergence of the peak of the distribution, which they evaluate retrospectively on sessions of 150 stimuli. Specifically, they assume that the probability of detection is a modified sigmoidal function of the electric field magnitude whose threshold depends on the in-plane direction of the field. Their approach has several potential shortcomings: (1) binarizing the response discards differences in MEP amplitude above and below that threshold, and hence loses potentially valuable information; (2) the assumption of a sigmoidal detection probability function is heuristic and not rigorously grounded; (3) the full posterior of that approach is costly to compute and therefore not available in real time; (4) in consequence, the stopping rule has to rely on convergence alone and cannot use any quantitative probability or confidence measure.

The BaMS3 method by Schultheiss et al. (2026) formulates motor mapping as spatially structured Bayesian inference. It models graded MEP responses with a Hill-type sigmoid function of electric-field magnitude and includes noise terms. This relationship remains hy-pothetical and follows conventions in previous literature. Full inference takes approximately 5.4 minutes per fit and is therefore unavailable during a real-time session.

These approaches motivate a simpler formulation that can use graded responses and update location uncertainty between pulses. We seek three properties: (1) an interpretable probability map for defining stopping criteria, (2) an explicit assessment of convergence, and (3) computation within a typical interval between stimulations of about 5 seconds.

Graded MEP amplitudes provide information beyond whether a response crosses a detection threshold. However, the cortical electric field and the peripheral MEP are linked through several biophysical processes, making their exact functional relationship difficult to specify. We therefore assume a monotonic relationship, expressed as a linear association between field and MEP ranks, so that a stronger field evokes an MEP that is at least as large. This gives an exact quantitative hypothesis that holds with reasonable certainty over the stimulation strengths used and can be incorporated into probabilistic inference.

Here, we implement this scheme and test it against the three requirements (Fig. 1). Every element of the cortical region of interest is scored by Spearman’s rank correlation between the induced electric field and the MEP amplitude, a map that is recomputed within milliseconds after each pulse. An evidence function converts the score map into a posterior probability map over cortical location. The mass near the reported hotspot and the area holding a given fraction of that mass provide readouts of localization uncertainty. A stopping rule requires the local mass to remain above a preset confidence level, and we compare that level with the realized success rate.

**Figure 1:**
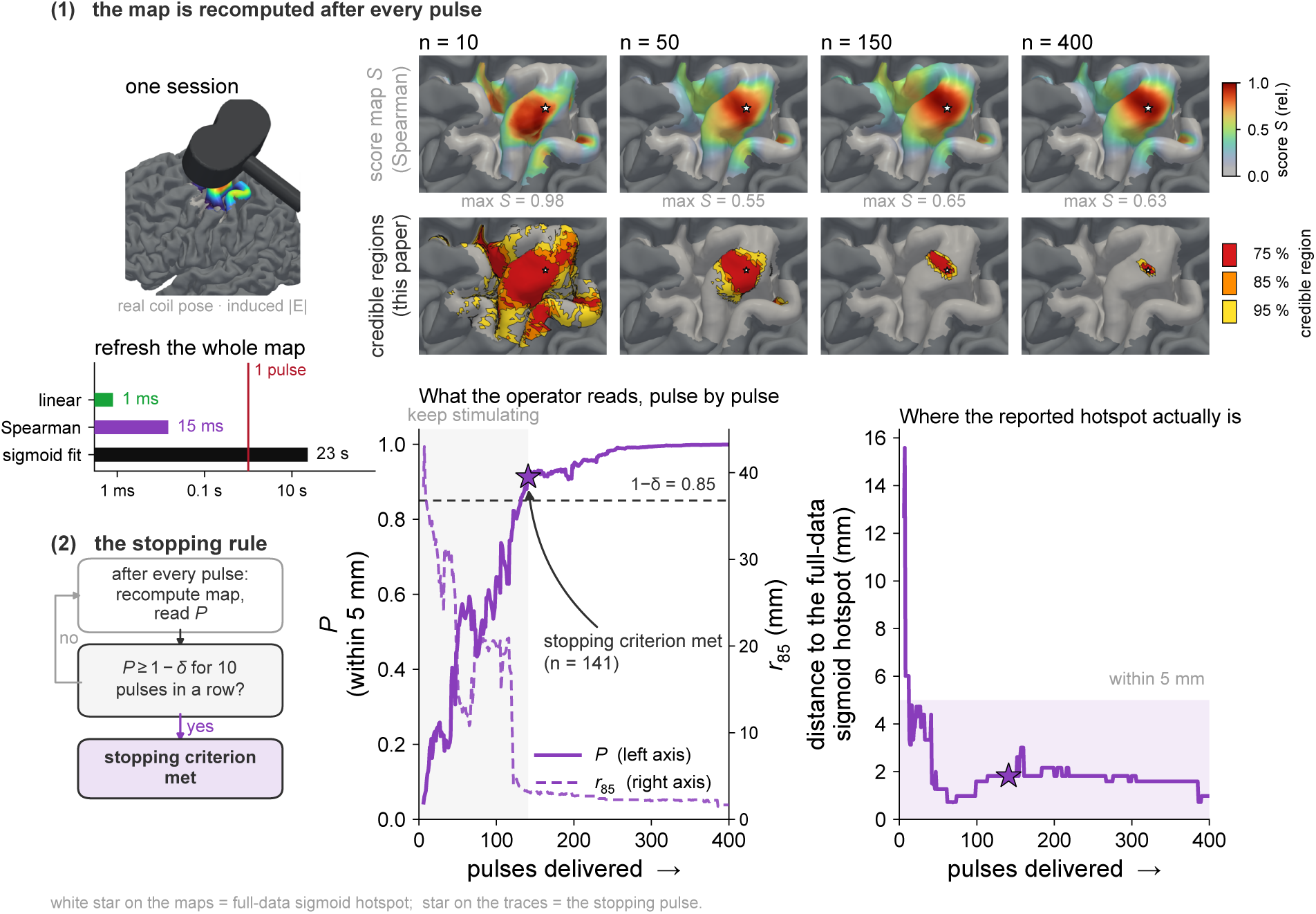
Overview of the method on one session. (cohort A, FDI, Spearman score). (1) After every pulse the score map *S* is recomputed (top row, 10 to 400 pulses) and converted into a posterior probability map, shown as credible regions holding 75%, 85% and 95% of the probability (second row); left, time to refresh the whole map. (2) Mapping stops once *P*_5_, the posterior probability within 5 mm of the hotspot, has stayed above 1 − *δ* = 0.85 for ten pulses (here at pulse 141). *r*_85_, radius around the hotspot holding 85% of the probability. Right, distance from the reported hotspot to the full-data sigmoid hotspot.

Independent functional validation is also needed. Same-subject comparisons have shown that hotspots obtained with conventional and other electric-field-based estimators lie within the same hand motor region as hand-movement fMRI (Chen et al., 2023; Diekhoff et al., 2011; Herwig et al., 2002). The nonlinear electric-field–MEP regression framework itself, however, has been evaluated primarily through within-TMS convergence and prospective testing of model-derived coil placements. It has not yet been evaluated against an indepen-dent functional measurement in the same individuals.

We evaluated the framework in two stages: (1) on calibrated virtual subjects with a known source, (2) on two independently acquired cohorts against the conventional full-data sigmoid hotspot, and, in one cohort, against the finger-tapping fMRI landmark of the same individuals.

## 2. Materials and methods

### 2.1. Participants and data

The two cohorts were acquired independently, with different coils, neuronavigation systems and coil-placement procedures (Fig. 2).

**Figure 2:**
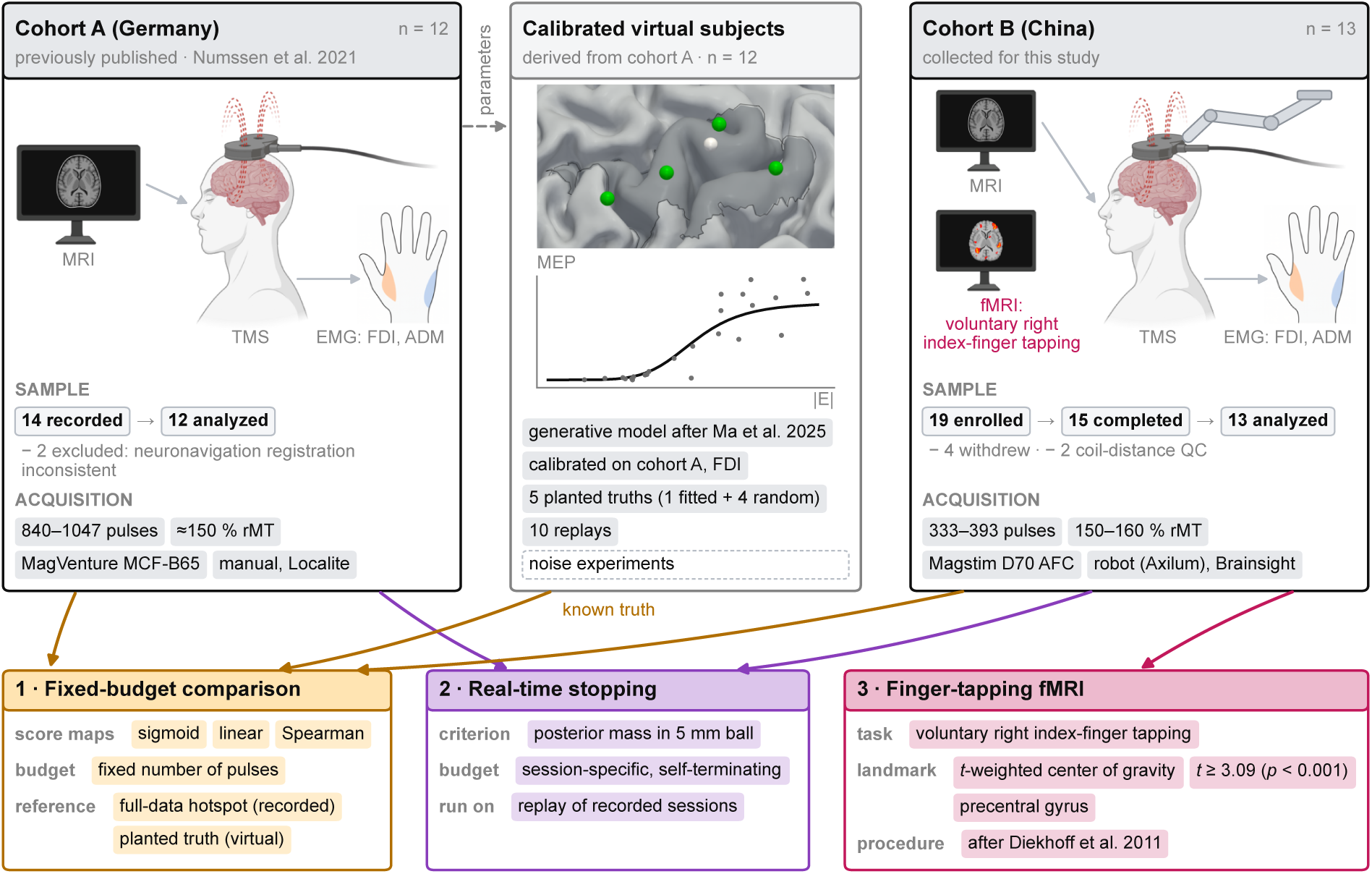
Datasets and analyses. Top row, the three datasets: cohort A (Germany, previously published; Numssen et al., 2021), virtual subjects calibrated on its FDI sessions (generative models after Ma et al., 2025, with the source element planted and known; Appendix D), and cohort B (China, collected for this study). Bottom row, the three analyses. Arrows are colored by the analysis they enter.

Cohort A was recorded at the Max Planck Institute for Human Cognitive and Brain Sciences following the established protocol (Numssen et al., 2021; Weise, Numssen, et al., 2023). The data were previously published (Numssen et al., 2021). Fourteen sessions were recorded (seven females, aged 21–38 years). Two were excluded: their archived neuronavi-gation registrations disagreed by more than 10 mm in the coil-to-head transform during the sessions. The remaining 12 participants each received 840–1047 single pulses. The intensity was fixed at about 150% of the resting motor threshold (Numssen et al., 2021), and the pulses were delivered with a MagVenture MCF-B65 figure-of-eight coil moved manually to randomly varied placements over the left hand knob under Localite neuronavigation.

Cohort B was collected for this study at the Center for Cognition and Brain Disorders, Hangzhou Normal University. Nineteen volunteers were enrolled, four of whom withdrew consent before completing the mapping session. Two of the 15 completers were excluded after coil-to-scalp distance quality control removed the first 100–135 pulses of their sessions. The final sample included 13 healthy, right-handed adults (nine women; age range, 22 to 29 years). Single-pulse TMS was delivered at 150% to 160% of the resting motor threshold with a Magstim Rapid^2^ stimulator and a D70 AFC coil positioned by a robotic arm (Axilum Robotics) under Brainsight neuronavigation, with surface EMG recorded simultaneously from the right first dorsal interosseous (FDI) and abductor digiti minimi (ADM). After quality control, 333 to 393 pulses per participant remained for analysis. All 13 participants also completed right index-finger tapping fMRI (Section 2.7). Appendix C gives the full acquisition parameters.

FDI and ADM were analyzed separately. Each participant–muscle pair is one localization problem. For each participant, an individual SimNIBS 4.5 head model was built with charm (Puonti et al., 2020; Thielscher et al., 2015). The region of interest (ROI) was the mid-layer triangular mesh over the primary motor cortex (8 921–17 609 elements; median element area 0.30 mm^2^ in cohort A and 0.34 mm^2^ in cohort B). For every pulse, |*E*| was computed at every element and rescaled by that pulse’s recorded stimulator output.

### 2.2. Reference hotspot

For every problem, a four-parameter log-sigmoid of MEP on |*E*| was fitted at every element using all of that participant’s pulses, and the element with the highest *R*^2^ was taken as the reference hotspot (Numssen et al., 2021; Weise, Numssen, et al., 2023).

### 2.3. Real-time score maps

The simplest per-element score *S* ∈ [0, 1] is the squared Pearson correlation, *S* = *R*^2^, computed from a linear fit to the raw MEP amplitudes of the *n* pulses delivered so far. It has a closed-form solution and can be recomputed between pulses (Appendix A). However, a linear fit does not capture the full dose–response relationship. The MEP saturates with |*E*|, so response curvature caps the linear *R*^2^ even at the reference hotspot, and compresses the contrast between hotspot and background on which localization depends, so that a larger area of cortex scores as high as the source in our virtual experiment (Fig. 3).

**Figure 3:**
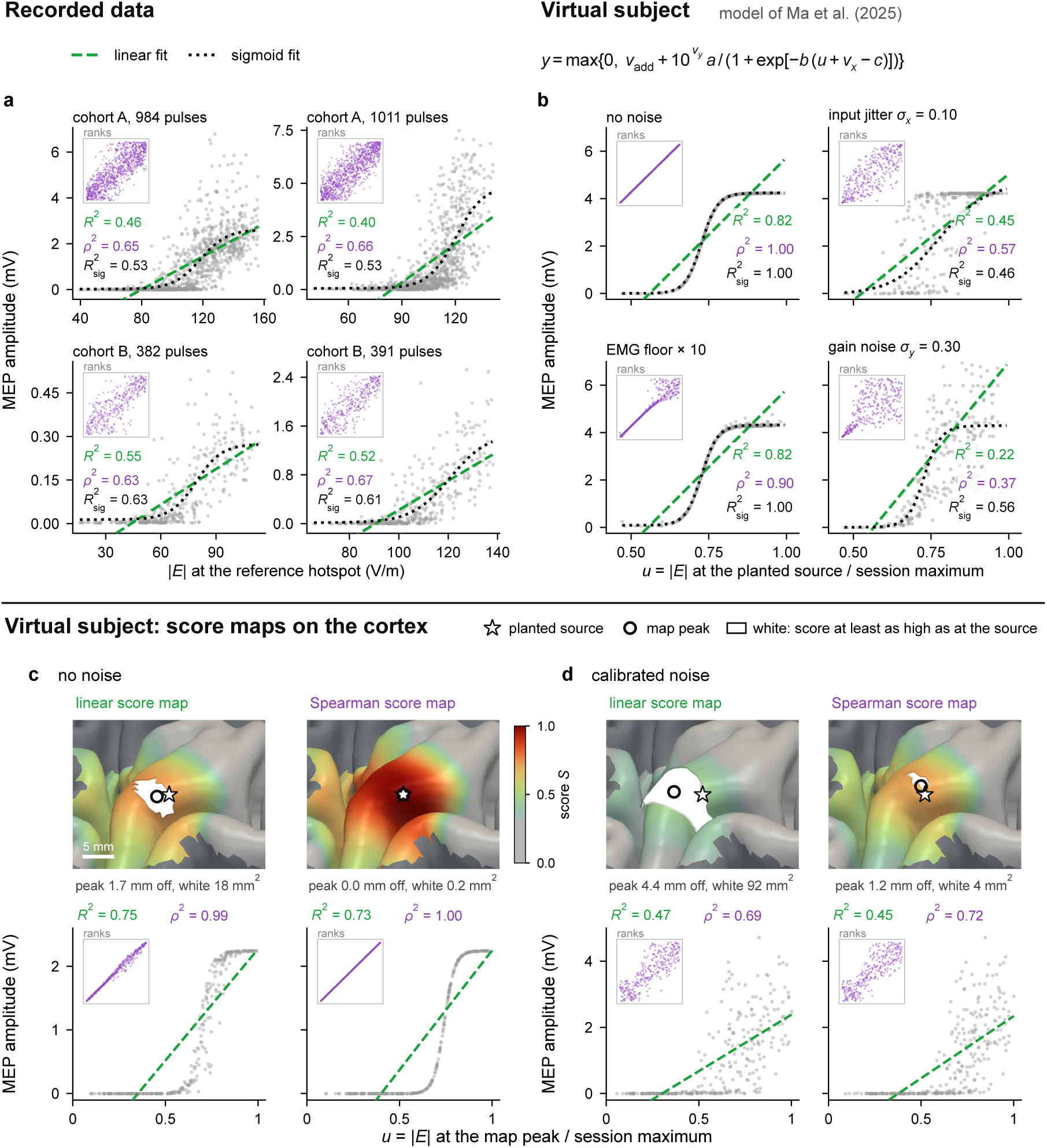
Saturation of the MEP input–output relation and its effect on the linear and Spearman scores. Dashed, linear fit; dotted, sigmoid fit; insets, the same data as ranks. Printed values: linear *R*^2^, Spearman *ρ*^2^ and the *R*^2^ of the sigmoid fit (*R*^2^). (a) Recorded MEPs against the electric field at the reference hotspot, two FDI participants per cohort. (b) A virtual subject at its planted source, without noise and with one source of variability at a time: input jitter (*σ_x_*), an elevated EMG floor and gain noise (*σ_y_*); model in Appendix D. (c, d) Linear and Spearman score maps of a second virtual subject after 300 pulses, without noise (c) and with all three sources at the levels calibrated to cohort A (d). White, cortex scoring at least as high as the planted source, i.e., the area the map cannot separate from it. Below each map, the MEPs at its peak. *u*, electric field divided by its session maximum.

The expected MEP increases with effective stimulation, but threshold, slope and sat-uration vary between participants (Devanne et al., 1997; Numssen et al., 2024) and with the local field orientation (Weise, Worbs, et al., 2023); what these relationships share is monotonicity. Spearman’s *ρ*, the Pearson correlation of ranks, measures monotonic associ-ation without specifying its form. Being invariant under increasing transformations, it is not reduced by saturation (Fig. 3) and limits the influence of extreme amplitudes. We take *S* = *ρ*^2^, analogous to the linear *R*^2^.

### 2.4. From a score map to a posterior probability map

The score summarizes the association between electric-field magnitude and MEP ampli-tude, but does not account for the number of pulses supporting that association. For the linear score, we incorporate sample size through the marginal likelihood, which averages the likelihood over the unknown regression parameters and noise scale (Kass & Raftery, 1995). Under the Gaussian linear model and priors specified in Appendix B, including the Jeffreys prior on the noise scale (Jeffreys, 1946), this integration yields the evidence

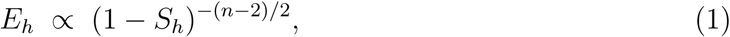

where *S_h_* is the linear score at element *h* and *n* is the number of pulses. The derivation is given in Appendix B, Eq. (B.2). With this exponent, the same association receives greater weight when supported by more pulses.

The Spearman score measures agreement between ranks and requires a different evidence function. We assess this agreement against what would be expected if fields and MEPs were randomly paired. The resulting Bayes factor is defined on the rank correlation itself (Johnson, 2005; Yuan & Johnson, 2008), using a normal approximation with null variance 1*/*(*n* − 1) (Hotelling & Pabst, 1936). It gives *E_h_* ∝ exp (*n* − 1)*S_h_/*2 (Appendix B).

For either score, normalizing *E_h_* over the ROI gives the posterior probability map, *p_h_* = *E_h_/ _j_ E_j_*, under the single-source model described in Appendix B. The prior is uniform over the elements of the ROI. Over the cortical surface we display nested credible regions: the smallest sets of elements that hold 75%, 85% and 95% of the posterior mass, obtained by the highest-posterior-density ranking defined in Section 2.5.

### 2.5. Stopping criterion and resolution readout

For the stopping rule, the reported hotspot is the element whose surrounding 5 mm geodesic ball contains the largest posterior mass. We summarize the posterior using two measures. *P*_5_ is the posterior mass within a geodesic ball of 5 mm around the currently reported hotspot, and the session may stop once *P*_5_ has held at or above 1 − *δ* for ten consecutive pulses. *A*_85_ is the area of the highest-posterior-density set containing 85% of the posterior mass. We rank elements by posterior probability per unit area, *p_h_/a_h_*, where *a_h_* is the area of element *h*. We then select elements in descending order until their probabilities sum to 0.85 and add their areas (Fig. 1).

### 2.6. Evaluation

We evaluated convergence, stopping performance, and spatial resolution. Convergence was quantified as the geodesic distance between the score-map peak after the first *n* pulses and a reference location. For recorded sessions, where the true source is unknown, the full-data sigmoid hotspot served as the reference, thereby measuring agree-ment with the established estimator. Localization accuracy against a known source was additionally assessed in 12 virtual subjects, one per cohort A participant. Each generative model was calibrated to the corresponding session, with five planted source locations per subject (Appendix D). Convergence was then measured relative to the planted source.

Stopping performance was evaluated on recorded sessions replayed in random pulse or-ders, using the posterior-based hotspot. We report the stopping pulse count, geodesic error relative to the full-data sigmoid reference, and the proportion of stopped replays within 5 mm of that reference, compared with the nominal level 1 − *δ*. Spatial resolution was summarized by *A*_85_.

### 2.7. Cross-modal comparison with finger-tapping fMRI

The 13 participants of cohort B also performed a right index-finger tapping task, a movement driven mainly by the FDI, which is therefore the muscle used for the fMRI comparison. The primary fMRI landmark reproduced the procedure of Diekhoff et al. (2011): a *t*-weighted center of gravity of all supra-threshold locations in the participant’s precentral gyrus (*t* ≥ 3.09, one-tailed *p <* 0.001 uncorrected). We fitted the general linear model on each participant’s native cortical surface and smoothed along the cortical sheet (Brodoehl et al., 2020; Coalson et al., 2018), placing the fMRI and TMS estimates on the same individual mesh. Three-dimensional Euclidean distance was the primary cross-modal measure.

The primary landmark was compared with the full-data sigmoid, linear and Spearman FDI hotspots. To describe their relative cortical positions, we expressed the fMRI center of gravity and each TMS hotspot in a common anatomical frame: signed geodesic distance from the central-sulcus boundary, and position along the principal axis of that boundary relative to the MNI hand-knob coordinate mapped to each participant. The analysis was repeated in a combined precentral–postcentral mask as a sensitivity check. Acquisition and analysis details are given in Appendix C.

## 3. Results

### 3.1. Computation time and convergence of the real-time score maps

Updating the full ROI took 1.1–333 ms for the Spearman score and 0.14–33 ms for the linear score as the session grew from 10 to 1000 pulses, against 13.0–31.9 s for the sigmoid fit (Fig. 1). Extrapolated to a whole hemisphere, the sigmoid fit would take about nine minutes, against 1.6 s and 224 ms for the Spearman and the linear score, respectively (Appendix A). In the recorded-data replays, Spearman and sigmoid generally showed smaller mean distances to the full-data reference and less dispersion than the linear score (Fig. 4a). Their performance was broadly comparable, with Spearman showing lower mean distance and dispersion in cohort A FDI.

**Figure 4:**
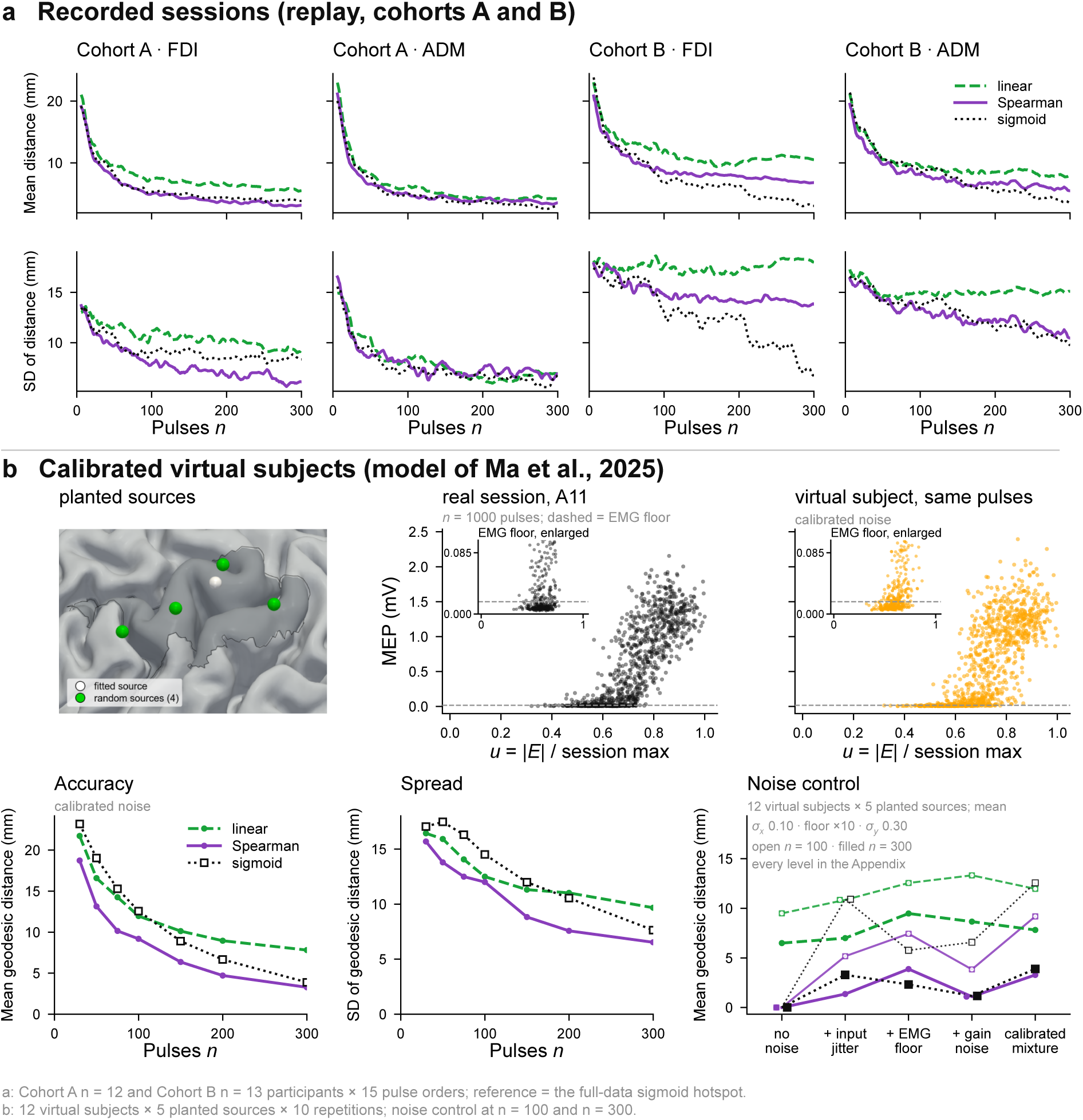
Convergence on recorded sessions and on calibrated virtual subjects. (a) Recorded sessions: mean (top) and standard deviation (bottom) of the geodesic distance from the map peak after *n* pulses to the full-data sigmoid hotspot, per cohort and muscle. Dotted, the sigmoid fitted to the same *n* pulses. (b) Calibrated virtual subjects. Top: planted sources on one subject’s cortex, and MEPs against the electric field in the recorded session and in its virtual subject. Bottom: mean and standard deviation of the geodesic distance to the planted source, and (right) the error without noise, with each source of variability alone and with all three (Appendix D).

Across the 12 virtual subjects and five planted sources per subject, Spearman showed lower mean localization error and dispersion than sigmoid at the tested pulse counts (Fig. 4b). The difference in median error was larger at 50 and 100 pulses and small at 300 pulses (Ta-ble D.1). The noise-control experiments showed that Spearman was less affected by input jitter than sigmoid, although an additive EMG floor disrupted the ordering of small responses (Fig. 3b; Appendix D). When the stimulator output was reduced so that fewer pulses evoked an MEP, all three methods lost accuracy, the sigmoid the most, and the Spearman score remained the most accurate (Fig. D.3).

### 3.2. Concentration of the posterior with the number of pulses

The score map alone did not indicate when enough pulses had been delivered. In the session of Fig. 1, the highest linear score in the ROI was 0.61 after 10 pulses and 0.44 after 400, because with few pulses high scores arise by chance. Across participants, the linear score at the reference hotspot ranged from 0.16 to 0.72, so no single score value could serve as a stopping threshold.

The posterior probability map concentrated as pulses accumulated (Fig. 5). *A*_85_ shrank between 10 and 300 pulses in all 50 localization problems. In cohort A, its median fell from 1702 to 4.9 mm^2^ with the linear score and from 1988 to 7.3 mm^2^ with the Spearman score; in cohort B, from 928 to 2.7 mm^2^ and from 1771 to 5.2 mm^2^. At 300 pulses, the Spearman region was larger than the linear region in 38 of 50 problems. Section 3.3 tests whether these concentrations match the realized accuracy.

**Figure 5:**
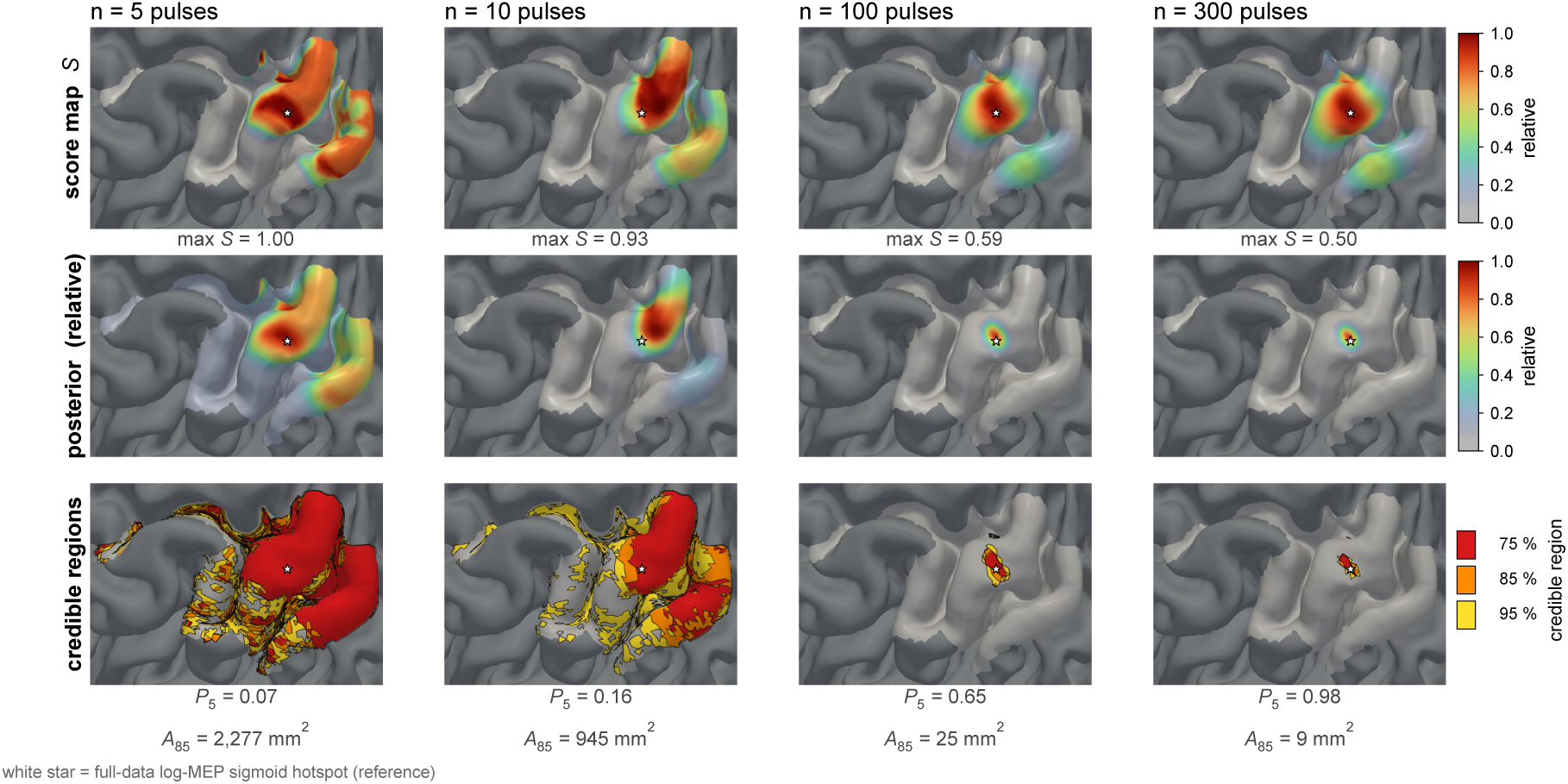
From score map to posterior probability map. (one cohort A participant, FDI, Spearman score) at 5, 10, 100 and 300 pulses. Rows: the score map *S*; the posterior probability, which combines *S* with the number of pulses (relative scale); and the same posterior as credible regions holding 75%, 85% and 95% of the probability.

### 3.3. Stopping performance and per-session pulse requirements

Across both cohorts, at 1−*δ* = 0.85 the stopping criterion was met in 96–100% of replays, each localization problem met the criterion in at least one replay, and the hotspot at stopping lay a median of 1.6–1.9 mm (Spearman) and 2.0–2.4 mm (linear) from the full-data sigmoid hotspot (Table 1).

**Table 1:**
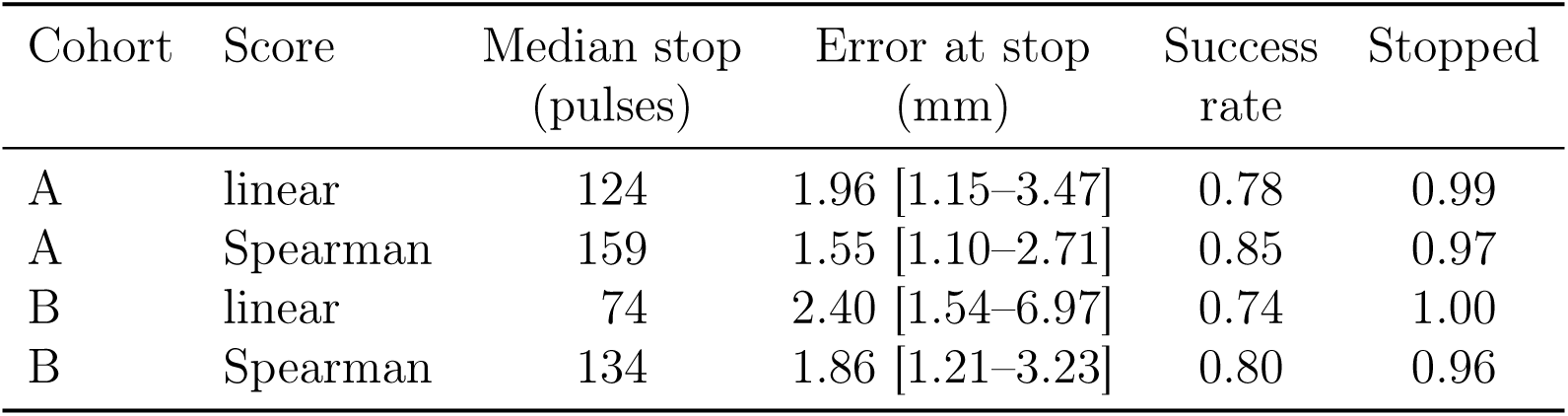
Stopping criterion at 1 − *δ* = 0.85, held for ten pulses, over replays of each session in random pulse order, up to its full pulse count. Error at stop, geodesic distance from the hotspot at stopping to the full-data sigmoid hotspot, median [IQR] over replays that stopped; success rate, fraction of these replays with an error of 5 mm or less; stopped, fraction of all replays that met the criterion.

At 1 − *δ* = 0.85, the Spearman score met the stopping criterion later than the linear score and reached a higher success rate, 0.85 against 0.78 in cohort A and 0.80 against 0.74 in cohort B (Table 1). Sweeping 1 − *δ* from 0.5 to 0.95 (Fig. 6a), the Spearman hotspot at stopping moved closer to the full-data sigmoid hotspot and remained closer than the linear hotspot at every matched budget. The Spearman success rate rose with 1 − *δ* in both cohorts, from 0.78 to 0.89 in cohort A and from 0.70 to 0.83 in cohort B, and was higher than the linear success rate at all four levels. Both scores met the nominal level at 0.5 and 0.7 in both cohorts. At 0.85 only the Spearman score in cohort A met it, and at 0.95 neither score did.

**Figure 6:**
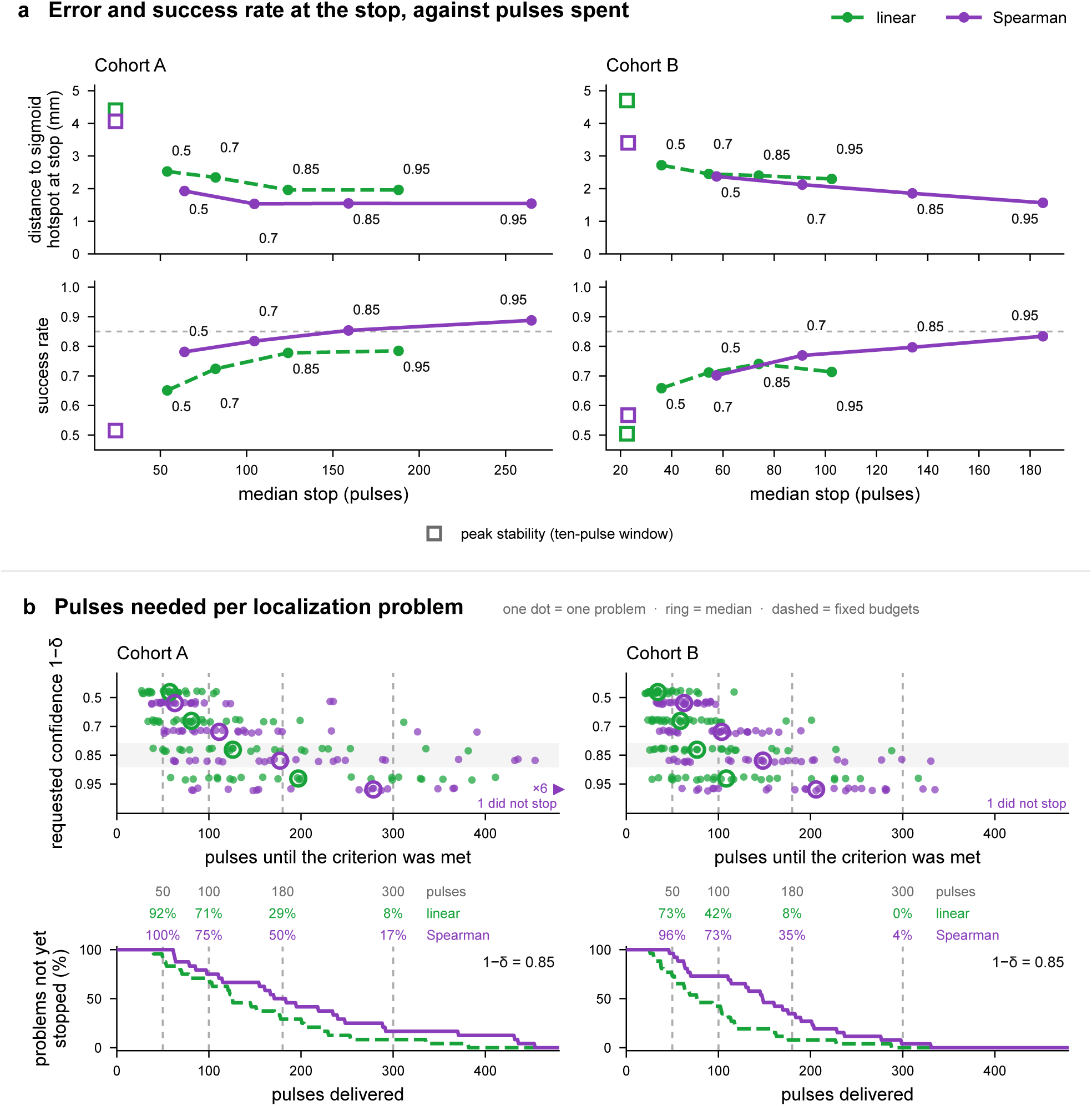
The stopping criterion. Cohort A left, cohort B right. (a) Error at the stop (top) and success rate (bottom) against the median number of pulses at the stop, for requested confidence 1 − *δ* from 0.5 to 0.95 (labels); squares, stopping once the reported hotspot has stayed within 5 mm for ten pulses. (b) Top: pulses until the criterion was met, one dot per localization problem (participant and muscle; median over eight pulse orders); dashed, fixed budgets from the literature. Bottom: fraction of problems still mapping at 1 − *δ* = 0.85.

With the same posteriors, a peak-stability rule (stop once the reported hotspot has stayed within 5 mm for ten pulses) stopped after about 25 pulses at a success rate near 0.5 and fell below a success rate of 0.85 in most problems (open squares in Fig. 6a).

The number of pulses required by individual localization problems varied by an order of magnitude (Fig. 6b). The Spearman medians, 134 and 159 pulses, are close to the approxi-mately 180 pulses estimated retrospectively as the minimum for a stable map (Numssen et al., 2021). Applied to an individual session, however, a fixed budget could exceed or fall short of the pulse count required by the stopping criterion. At 180 pulses, at worst half of problems had still not reached the requested confidence, while the fastest problems in every cohort–score combination had reached it within 70 pulses.

### 3.4. Cross-modal comparison with finger-tapping fMRI

The primary fMRI landmark was defined in all 13 participants. Figure 7a shows the FDI hotspots and finger-tapping activation on each participant’s cortical surface. Across the three TMS estimators, the Euclidean distance of the landmark from the FDI hotspots was 8.3 ± 4.3 mm (mean ± SD; median 8.0 mm; 39 participant–method comparisons). This distance lies within the range reported in same-subject comparisons using other TMS map-ping approaches: approximately 10 mm (Herwig et al., 2002), 10.5 ± 3.0 mm (Diekhoff et al., 2011) and 20.5 ± 8.5 mm (Chen et al., 2023).

**Figure 7:**
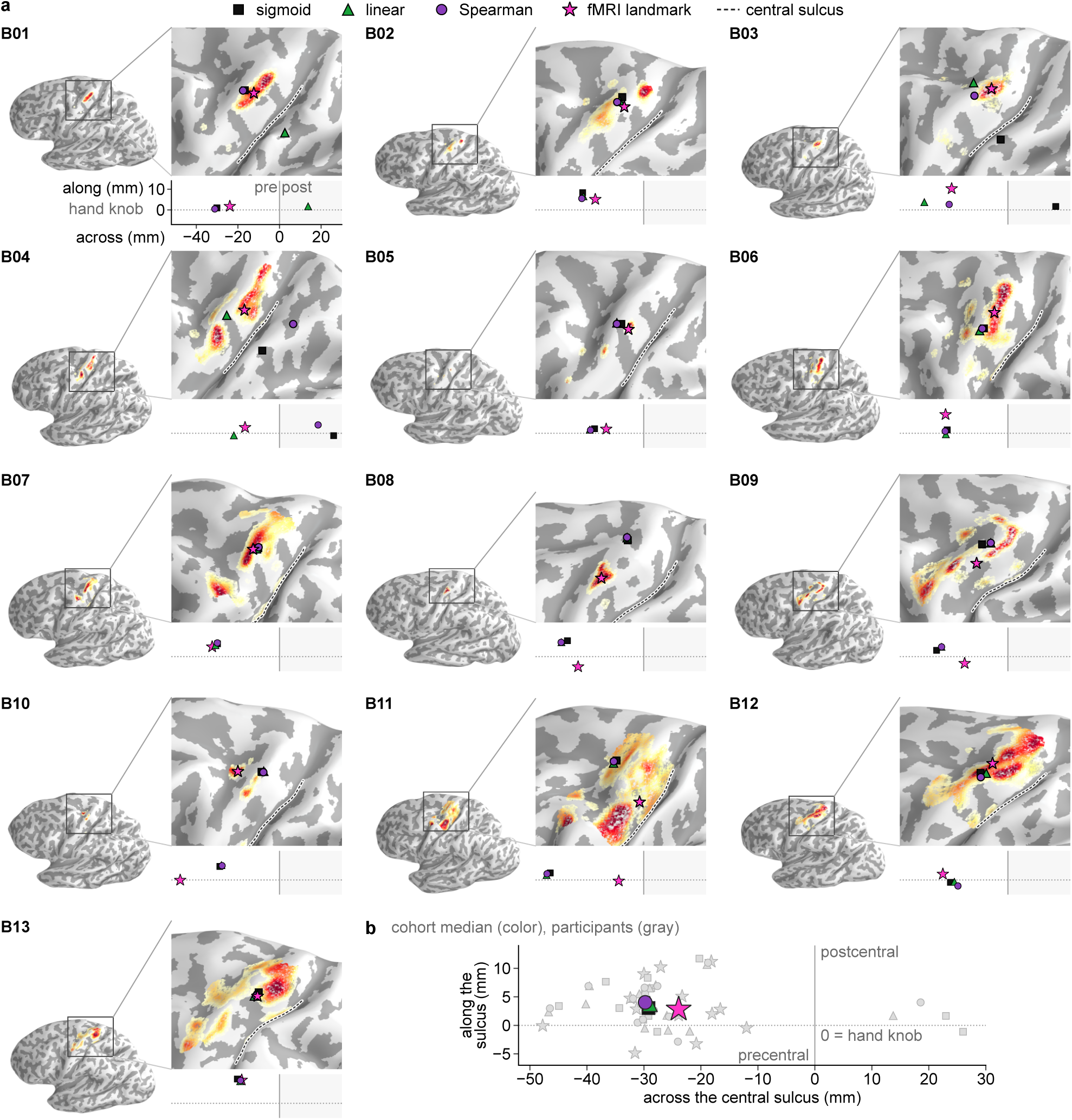
FDI hotspots and the fMRI landmark. (cohort B, *n* = 13). (a) One panel per participant. Left, inflated left hemisphere; the box marks the region enlarged on the right. Gray shading, curvature (dark, sulcus); dashed line, central sulcus; color, tapping activation from *t* ≥ 2.3 (*p <* 0.01, display); the landmark is the *t*-weighted center of gravity of its *t* ≥ 3.09 (*p <* 0.001) part (one-tailed, uncorrected). Below the enlarged view, the same locations in a common anatomical frame, with axes labeled for B01. Across: signed geodesic distance to the precentral–postcentral boundary (negative, precentral; shaded, postcentral). Along: position on the principal axis of that boundary relative to the MNI hand-knob coordinate mapped to each participant (positive, superior–medial). (b) All participants in the same frame (gray) and cohort medians (color).

In the common central-sulcus frame, the cohort medians of the fMRI and TMS locations lay on the precentral bank near the hand knob (Fig. 7b). Relative to the fMRI center of gravity, the paired median TMS coordinates lay 1.2–5.4 mm farther onto the precentral bank for the three estimators, although none of these differences was significant. When the fMRI search mask was expanded from the precentral gyrus to include the postcentral gyrus, the median fMRI coordinate moved toward the opposite bank from −23.9 mm to −12.3 mm, and 5 of 13 centers of gravity lay postcentrally.

## 4. Discussion

In this study, we showed that a rank-correlation score map can be recomputed between pulses and converted into a posterior probability map that defines a stopping criterion for each session. On the recorded sessions, the Spearman hotspot and a sigmoid fitted on the same pulses were at broadly comparable distances from the full-data sigmoid reference, and on virtual subjects with a planted source the Spearman score had a lower mean localization error than the sigmoid fit. The number of pulses needed to meet the stopping criterion varied by an order of magnitude between participants and muscles.

Rank correlation is a practical substitute for repeated nonlinear fitting. The saturation that caps the linear score (Fig. 3) does not reduce it in the noise-free limit. In the noise controls, input jitter degraded the Spearman score less than the sigmoid fit. Variability in the effective stimulation is amplified along the steep part of the recruitment curve, and ranks are insensitive to the resulting changes in response magnitude (Appendix D). An additive EMG floor obscures the ordering of small responses and limits the Spearman score; in the virtual subjects, this effect was limited (Appendix D).

Pulse requirements differed by an order of magnitude between sessions, so a fixed budget exceeds the requirement of some sessions and falls short of others. Posterior probability mass within a fixed radius quantifies confidence at stopping. The requested confidence sets the balance between pulse count and accuracy; for the Spearman score, raising 1 − *δ* delayed the stop and increased the success rate in both cohorts. The linear criterion stopped earlier but was overconfident, its success rate falling further below the requested level, which is undesirable in clinical use; the Spearman criterion required more pulses and was more accurate at stopping.

The method extends the electric-field-informed regression framework by changing the score and its uncertainty readout (Fig. 8). Its only assumption about the response is that the MEP increases monotonically with the field; no recruitment curve is fitted. On this basis, the posterior gives each cortical element a probability of being the source, so the stopping criterion is a statement about location whose realized success rate can be checked against the requested level. Convergence is read from the same posterior after every pulse, as the confidence in the current hotspot (*P*_5_) and the cortical area that remains plausible (*A*_85_). The Spearman map was updated within a third of a second, well within the interval between pulses. The two Bayesian formulations model the response explicitly, as a sigmoidal detection probability of binarized responses (Laine et al., 2026) or as a Hill-type sigmoid of graded MEPs (Schultheiss et al., 2026), and their full posteriors take minutes to compute. The rank score uses the graded amplitudes without such a model.

**Figure 8:**
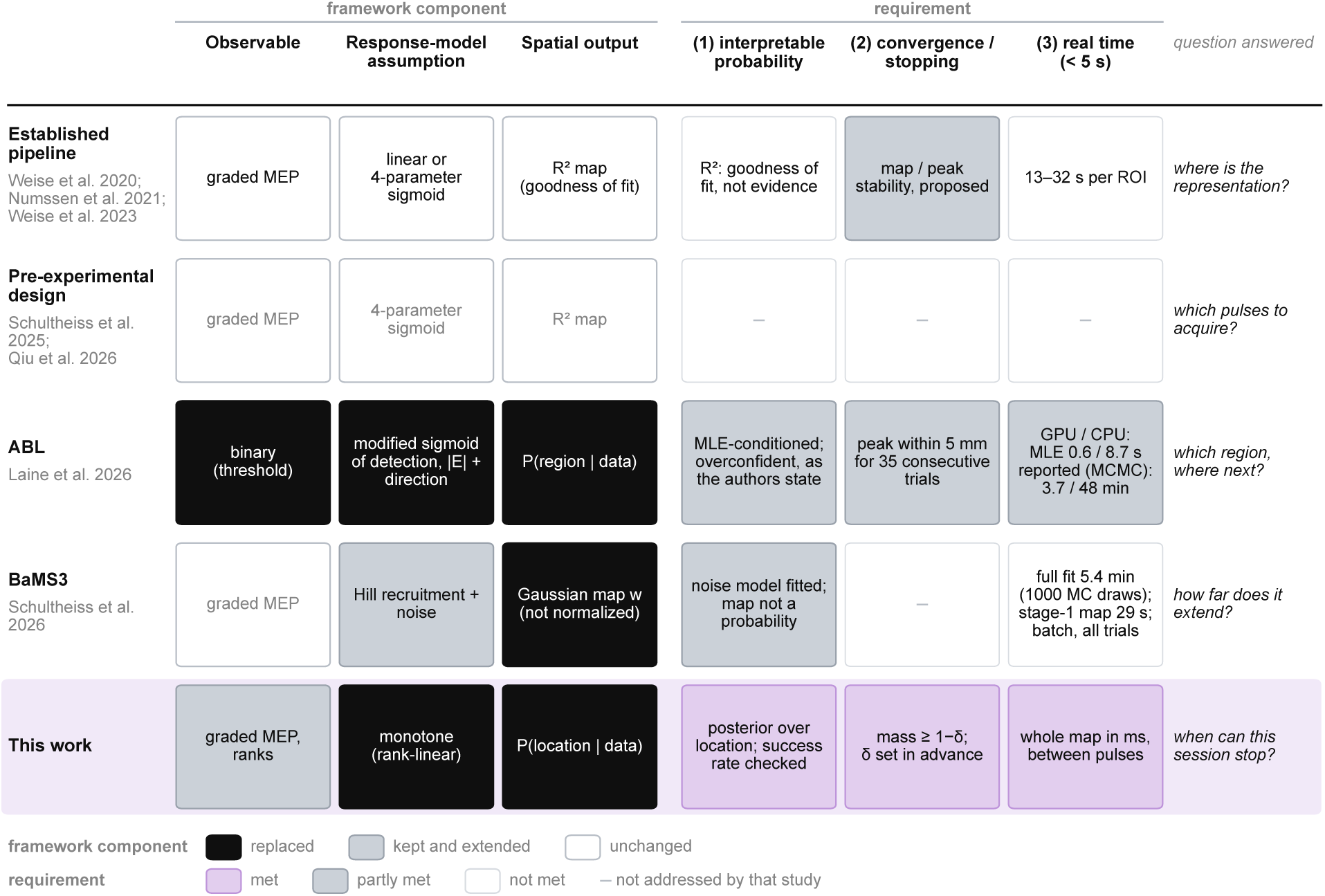
Lineage of electric-field-informed motor mapping. Each row is a branch of the same framework. Left: which framework component it changes relative to the established pipeline (Numssen et al., 2021; Weise, Numssen, et al., 2023; Weise et al., 2020), the observable, the assumed response model and the spatial output. Right: what each branch provides for the three requirements stated in the Introduction: (1) a quantitatively interpretable probability of cortical location, (2) convergence or stopping and (3) computation possible within the interval between pulses. Computation times are those reported by each study. MLE, maximum-likelihood estimate; MC, Monte Carlo; MCMC, Markov chain Monte Carlo.

The FDI hotspots and the finger-tapping landmark, a functional reference independent of TMS, lay in the same precentral hand region, at distances within the range of earlier same-subject comparisons (Chen et al., 2023; Diekhoff et al., 2011; Herwig et al., 2002). The TMS hotspots tended to lie farther onto the precentral bank, and including the post-central gyrus in the fMRI search mask moved the landmark toward the postcentral bank. The two modalities localize the same hand region but measure different physiological sig-nals: finger tapping engages muscles other than the FDI, its sensory feedback activates the primary somatosensory cortex, and the loci of maximal BOLD response and maximal MEP elicitability need not coincide.

Several limitations remain. First, the stopping rule was evaluated on randomly permuted recordings, and its performance in the original acquisition order and in prospective sessions remains to be tested. Second, both cohorts comprised healthy adults, two intrinsic hand muscles and figure-of-eight coils. Applicability to other muscles and clinical populations is unknown. Patients with high motor thresholds may receive fewer pulses that evoke an MEP; in virtual subjects, a reduced stimulator output lowered the accuracy of all three methods (Fig. D.3). Third, the rank score assumes that the response increases monotonically with the field. This holds for muscle mapping but may fail for other regions or tasks, for example with U-shaped relations. Fourth, the posterior is conditional on the response model, the prior and the cortical ROI. Field direction can be added to the response model (Laine et al., 2026; Weise, Worbs, et al., 2023), and a spatially extended source model would allow neighboring elements to share information (Schultheiss et al., 2026). A prospective study of localization and success at stopping is the next step, followed by adaptive selection of the next coil placement.

## Ethics

Both studies were approved by the responsible ethics committees: cohort A by the Ethics Committee of the Medical Faculty of the University of Leipzig (Numssen et al., 2021) and cohort B by the Ethics Committee of the Center for Cognition and Brain Disorders, Hangzhou Normal University (No. 2021071002). All participants gave written informed consent, and all procedures conformed to the Declaration of Helsinki and to the safety guidelines for transcranial magnetic stimulation (Rossi et al., 2021).

## Declaration of generative AI and AI-assisted technologies in the manuscript preparation process

AI played no role in planning or carrying out the research described in this paper. AI (GPT-6 and Claude Opus 5) was used in writing the manuscript, to check spelling and grammar and to reword draft text. All AI-assisted text was verified and edited by the authors, who take full responsibility for the content of this paper.

## Author Contributions

Conceptualization: RQ, KW, and TRK; Methodology: RQ, KW, and TRK; Software: RQ and ZF; Formal analysis: RQ; Investigation: RQ, ZF, KW, and TRK; Data curation: ON, ZF and YZ; Resources: YZ, FW, KW, and TRK; Visualization: RQ; Supervision: KW and TRK; Project administration: TRK; Funding acquisition: YZ, FW, KW and TRK; Writing – original draft: RQ and ZF; Writing – review and editing: all authors. All authors discussed the results and approved the final manuscript.

## Data availability

The cohort A data were previously published. The electric fields within the ROI and the MEP data are available at https://osf.io/jygu2/ (Numssen et al., 2021). The cohort B data are not publicly available owing to privacy restrictions. They are available from the corresponding authors upon reasonable request. The code implementing the score maps, evidence functions and stopping criterion is available at https://github.com/HughYau/real-time-probabilistic-TMS-motor-mapping

## Appendix A. Update cost of the score maps

### A.1 Algorithms and complexity

For *n* pulses and *V* ROI elements, the linear score at every element follows from five sums over pulses: of the field, the squared field, the MEP, the squared MEP and the product of field and MEP. Computing them for the whole ROI is one pass over the *n* × *V* field matrix, with cost *O*(*nV*) and no loop over elements. A new pulse adds one term to each sum, so an incremental update costs *O*(*V*), independent of the session length. The Spearman score applies the same computation to ranks. Because a new pulse can change the ranks of earlier pulses, the ranks are recomputed by sorting the field values at each element, which costs *O*(*nV* log *n*). The sigmoid has no closed form. Its four parameters (midpoint, slope, amplitude and baseline) are fitted at every element by iterative nonlinear least squares with the settings of the established pipeline (Numssen et al., 2021; Weise, Numssen, et al., 2023), with up to 1000 function evaluations per element, and the fit is repeated after every new pulse.

### A.2 Timing measurements

Times were measured on one workstation (AMD Ryzen 9 9950X, 16 cores) with one recorded session (13 069 elements, 984 pulses) and refer to a full recomputation of the map (Table A.1). Larger maps were built by repeating elements and longer sessions by resampling pulses. The sigmoid ran on four parallel processes. At 300 pulses it took 28.7 s for this ROI and 205 s for 100 000 elements; extrapolated to one hemisphere (254 180 elements in the participant of Fig. 1), it would take 521 s.

## Appendix B. The two evidence functions

At each element *h*, both evidence functions quantify support for *h* as the source of the recorded responses relative to no association. The posterior map treats the elements of the ROI as mutually exclusive hypotheses for a single source with equal prior probability and normalizes the evidence, *p_h_* = *E_h_/ _j_ E_j_*. It is therefore conditional on the source lying inside the ROI and on the response model stated below. Both models take the pulses as independent draws.

**Table A.1.** : Time to refresh the whole map (seconds).

| *n* (at 13 069 elements) | 10 | 50 | 100 | 200 | 300 | 600 | 1000 |
|---|---|---|---|---|---|---|---|
| linear | 0.0001 | 0.0004 | 0.0008 | 0.0025 | 0.0043 | 0.0201 | 0.0331 |
| Spearman | 0.0011 | 0.0053 | 0.0147 | 0.0391 | 0.0602 | 0.1861 | 0.3331 |
| sigmoid | 13.0 | 14.5 | 23.0 | 25.6 | 26.2 | 25.3 | 31.9 |
| Elements (at 300 pulses) | 1 000 | 2 500 | 5 000 | 13 069 | 30 000 | 60 000 | 100 000 |
| linear | 0.0002 | 0.0008 | 0.0010 | 0.0049 | 0.0216 | 0.0465 | 0.0881 |
| Spearman | 0.0040 | 0.0101 | 0.0217 | 0.0654 | 0.1847 | 0.3753 | 0.6396 |
| sigmoid | 1.82 | 3.64 | 10.57 | 28.66 | 51.34 | 127.30 | 205.12 |

### B.1 Marginal likelihood of the linear score

At element *h* write the regression *y* = *a* + *b x_h_* + *ε* with *ε* ∼ N (0*, σ*^2^), where *x_h_* collects the pulse-wise field magnitudes at that element, standardized to unit variance across pulses, and *y* the responses. Placing flat priors on (*a, b*) and the Jeffreys prior *p*(*σ*) ∝ 1*/σ* on the noise scale (Jeffreys, 1946), the marginal likelihood (Kass & Raftery, 1995) is the integral of the Gaussian likelihood over the three nuisance parameters, which evaluates in closed form to

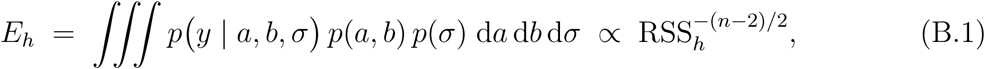

with RSS*_h_* the residual sum of squares of the fit at element *h*. The flat prior on *b* is improper and leaves an arbitrary constant in *E_h_*. This constant is the same at every element and cancels in the normalization. Standardizing *x_h_* leaves *R*^2^ unchanged and makes the design matrix, and with it the constant of the integral, the same at every element. Because RSS*_h_* = TSS (1 − *R*^2^) and the total sum of squares is shared by every element (the same responses enter every regression), TSS also cancels:

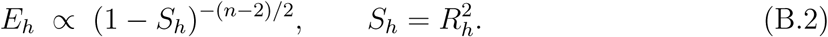

Elements whose fitted slope is negative receive *S_h_* = 0, the evidence value at zero associ-ation, because a response that falls as the field rises cannot mark the source. The model assumes Gaussian noise of constant variance on the raw amplitudes. Equation (B.2) is exact under that assumption and approximate for a saturating response with amplitude-dependent variability.

### B.2 Evidence for the rank score

A rank correlation has no Gaussian likelihood, so *ρ*^2^ cannot be substituted into Eq. (B.2). The pole at *S* = 1 would assign infinite evidence to a perfect ordering, an event of probability 2*/n*! under the null, and the two formulas diverge exactly at the high scores at which the stopping criterion is met. We follow the construction of Bayes factors on test statistics (Johnson, 2005), which Yuan and Johnson extended to nonparametric statistics with limiting normal distributions, including Spearman’s *ρ* (Yuan & Johnson, 2008). Under the null of no association Spearman’s *ρ* has mean zero, variance exactly *τ* ^2^ = 1*/*(*n* −1) (Hotelling & Pabst, 1936), and a distribution that is close to normal at the sample sizes used here. Writing

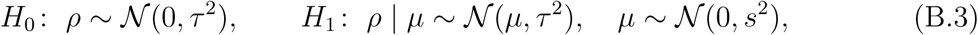

both marginal likelihoods are Gaussian and the Bayes factor is available in closed form,

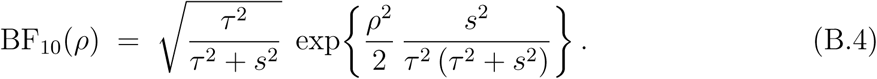

The prefactor does not depend on *ρ* and is common to all elements. We take the diffuse limit *s*^2^ → ∞, in which the exponent tends to *ρ*^2^*/*(2*τ* ^2^). Dropping the prefactor, as for the linear evidence, gives

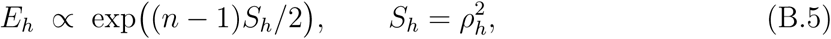

where, as in B.1, a negative rank correlation is assigned *S_h_* = 0. Two approximations are made. First, the normal law with the null variance is used for *ρ* under both hypotheses. Because *ρ* is bounded, its exact permutation distribution has lighter tails than the normal near |*ρ*| = 1, so Eq. (B.5) understates the evidence at high scores. Second, a Bayes factor on a statistic compares one element with the null of no association. Normalizing Bayes factors against a common null gives posterior model probabilities (Kass & Raftery, 1995). Because each *ρ_h_* is a different statistic of the data, this normalization is an approximation for the rank evidence (Johnson, 2005). For the linear evidence, a marginal likelihood of the data themselves, this step is exact.

## Appendix C. Cohort B: acquisition and fMRI protocol

### Participants

All 13 participants were classified as right-handed with the Chinese version of the Edinburgh Handedness Inventory (Yang et al., 2018).

### Imaging

GE SIGNA Architect 3 T. T1: inversion-prepared gradient echo, 1 mm isotropic, 256×256×192, TR/TE/TI 2678/2.98/1100 ms, flip angle 7°. T2: three-dimensional fast spin echo, 1 mm isotropic, TR/TE 3202/123 ms, two averages. Diffusion: 64 directions, 2 mm isotropic, TR/TE 10725/105.5 ms, with reversed phase encoding for distortion cor-rection. The mesh of the individual finite-element head model was locally refined within the predefined motor-mapping ROI. The cortical mid-layer triangular ROI contained 8 921–11 497 elements across participants.

### Stimulation

Single pulses were delivered with a Magstim Rapid^2^ stimulator and a D70 AFC coil (The Magstim Company Ltd, Whitland, UK), positioned by a robot (Axilum Robotics SAS, Schiltigheim, France) with a force sensor at the coil face under Brainsight 2.4.10 neuronavigation (Rogue Research Inc., Montreal, Canada). Stimulator output was set to 150–160% of each participant’s resting motor threshold, capped at 100% of maximum stimulator output (MSO). Actual output ranged from 91% to 100% MSO. The protocol comprised 200 coil poses across 153–169 scalp positions, with two pulses delivered 5 s apart at each pose. Sessions lasted 90–144 min, including breaks and repeat stimulation when required.

### Electromyography

Surface EMG was recorded simultaneously from the right FDI and ADM using a belly–tendon montage (Kleim et al., 2007) and the two-channel amplifier integrated into the neuronavigation system. Signals were sampled at 3 kHz over a ±100 ms window around the pulse (−50 ms to 150 ms in one participant). MEP amplitude was calculated as the peak-to-peak amplitude between 18 ms and 35 ms after the pulse, the same window as cohort A, converted from microvolts to millivolts so that both cohorts share one scale. Pulses delivered beyond the accepted coil-to-scalp distance were removed by the acquiring laboratory.

**fMRI.** Gradient-echo EPI on the same scanner: multiband factor 4, TR 0.7 s, TE 30 ms, flip angle 80°, 3 mm isotropic, 72 × 72 × 44, 600 volumes (420 s). Participants completed 80 trials. Each trial began with a fixation interval of 3.5, 4.2, 4.9 or 5.6 s, followed by a 0.7 s finger cue. Each interval occurred 20 times in pseudorandom order. Participants pressed a button with the right index finger when the cue appeared (Fig. C.1). The general linear model treated each trial as an instantaneous event and was fitted on each participant’s native cortical surface with a geodesic FWHM of 4 mm along the sheet (Brodoehl et al., 2020; Coalson et al., 2018).

**Figure C.1.**
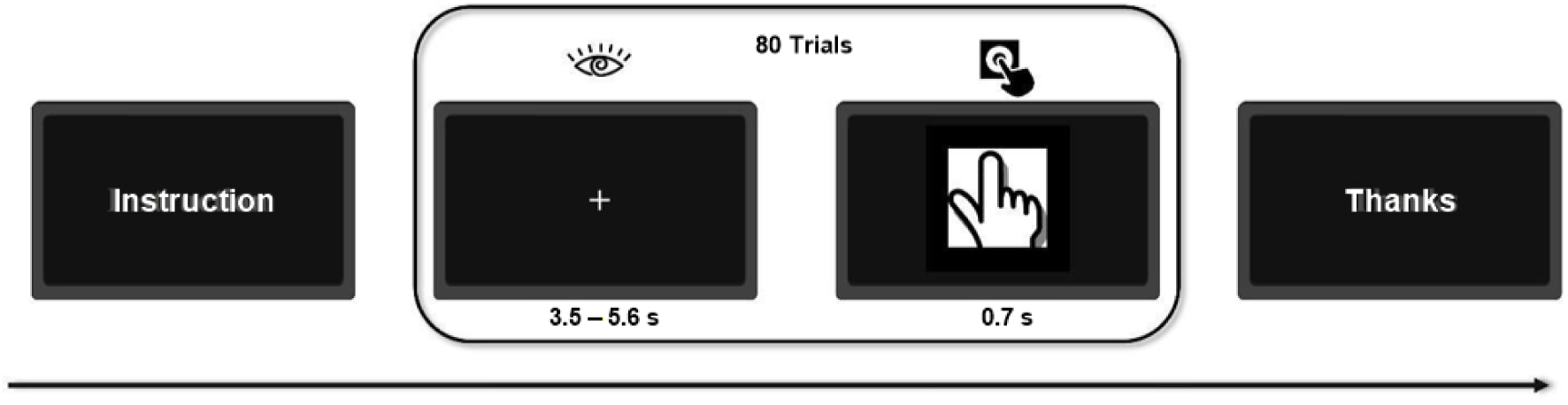
: **The finger-tapping task** (cohort B). Trial structure and timing; the jittered fixation interval preceded a finger cue, during which participants tapped a button.

## Appendix D. Calibrated virtual subjects

### D.1 Generative model

Each cohort A participant was paired with a virtual subject that generates MEPs from that participant’s own electric fields. For a planted source element *r*^∗^, the drive of pulse *i* is the normalized field magnitude at that element, *u_i_*= |*E_i_*(*r*^∗^)|*/* max*_j_* |*E_j_*(*r*^∗^)|. The response follows a log-logistic recruitment curve of the drive with no lower plateau and three variability sources (Ma et al., 2025): input jitter *v_x_* ∼ N (0*, σ*^2^) added to the drive before the nonlinearity, a multiplicative gain noise *v_y_* ∼ N (0*, σ*^2^) acting in the log_10_ domain, and an additive EMG floor *v*_add_ drawn from a generalized extreme value distribution,

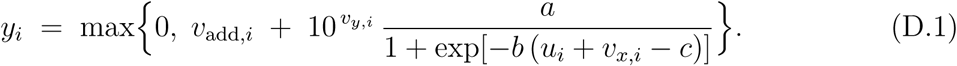

Its parameters were fitted per participant on that participant’s recorded session, and all 12 fits converged. Across the 12 virtual subjects, *σ_x_* was 0.052–0.149, *σ_y_* 0.051–0.289, and the floor location 0.0057–0.0175 mV, that is 0.2–1.8% of the saturation amplitude. Even without added noise, 27–81% of the pulses of a session fall below the 95th percentile of that floor.

### D.2 Planted sources

Five source elements were planted per subject, one fitted source and four random reach-able elements, resulting in 60 problems. An element *r* is reachable when the largest field the session delivers to it, relative to the largest field at the fitted source *r*_0_, reaches the midpoint of the recruitment curve, max*_i_* |*E_i_*(*r*)|*/* max*_j_* |*E_j_*(*r*_0_)| ≥ *c*.

### D.3 Conditions

The three variability sources were switched independently: the noise-free limit (all three set to zero); each source alone at a range of levels (*σ_x_*0.02–0.30, *σ_y_*0.05–0.50, floor scaled by a factor of 1 to 100); the calibrated mixture of all three; the mixture with one source removed; and the mixture with one source amplified.

### D.4 Evaluation

Within one condition, planted source and replay, the three score maps were computed on the same generated MEP sequence, so all comparisons are paired. The reported location is the argmax of the score map. We report the geodesic distance from that element to the planted source and the success rate, defined as a distance of 5 mm or less. The unit of analysis is a problem, i.e., one subject with one planted source. Under the calibrated mixture, all three score maps were replayed ten times, so the comparison there is paired replay by replay. In the remaining noisy conditions the sigmoid was replayed three times and the closed-form scores are restricted to those three paired replays. The noise-free limit is deterministic and was run once.

**Table D.1.** : Recovery of the planted source with all three sources of variability at their calibrated levels: median geodesic distance (mm) and success rate (distance of 5 mm or less) over 12 virtual subjects × 5 planted sou<u>rces × 10 repetitions.</u>

|  | *n* = 50 |  | *n* = 100 |  | *n* = 300 |  |
|---|---|---|---|---|---|---|
| Score | distance | success | distance | success | distance | success |
| sigmoid | 14.5 | 0.27 | 6.8 | 0.43 | 1.15 | 0.82 |
| Spearman | 7.8 | 0.39 | 4.1 | 0.54 | 1.06 | 0.85 |
| linear | 11.8 | 0.32 | 7.2 | 0.45 | 2.97 | 0.60 |

**Figure D.1.**
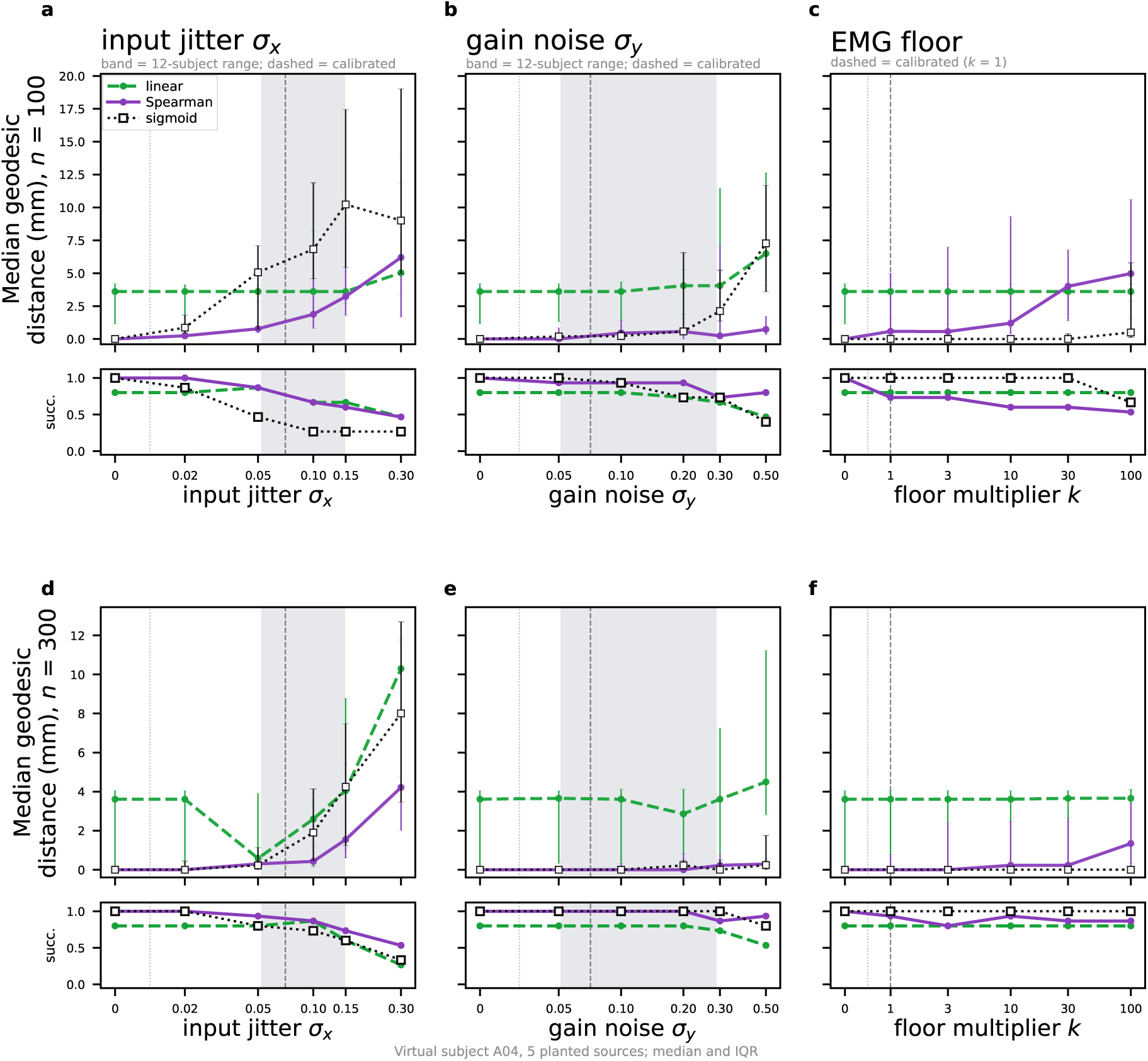
: **Effects of individual sources of variability.** Geodesic distance to the planted source against the level of input jitter, gain noise and EMG floor, with the other two set to zero, at 100 and 300 pulses; one virtual subject, five planted sources; the three scores are computed on the same simulated MEPs.

### D.5 Results

In the noise-free limit (zero level in Fig. D.1) the Spearman map and the sigmoid re-covered the planted source within the resolution of the mesh. The linear score retained a median error of 1.97 mm at 300 pulses. This residual error reflects model mismatch. The Pearson score is highest at neighboring elements whose field range produces a more linear field–MEP relationship. The linear peak lay more than 5 mm from the source in 20 of the 60 problems, and a median of 11.5 mm^2^ of cortex (interquartile range 2.2–59.1) scored at least as high as the source. For the Spearman score this set was the source element alone in all 60 problems. Fig. 3c shows one example.

Under the calibrated mixture, Spearman had the smaller median distance at low pulse counts and the difference narrowed as pulses accumulated (Fig. 4b, Table D.1). Across the 60 problems, the median paired difference in geodesic distance was −0.78 mm at 100 pulses (*p <* 0.001) and 0.00 mm at 300 pulses (*p* = 0.55; Fig. D.2). In the illustrated subject at 300 pulses, removing input jitter reduced the sigmoid median error to zero, while removing the floor reduced the Spearman median error to 0.23 mm.

**Figure D.2.**
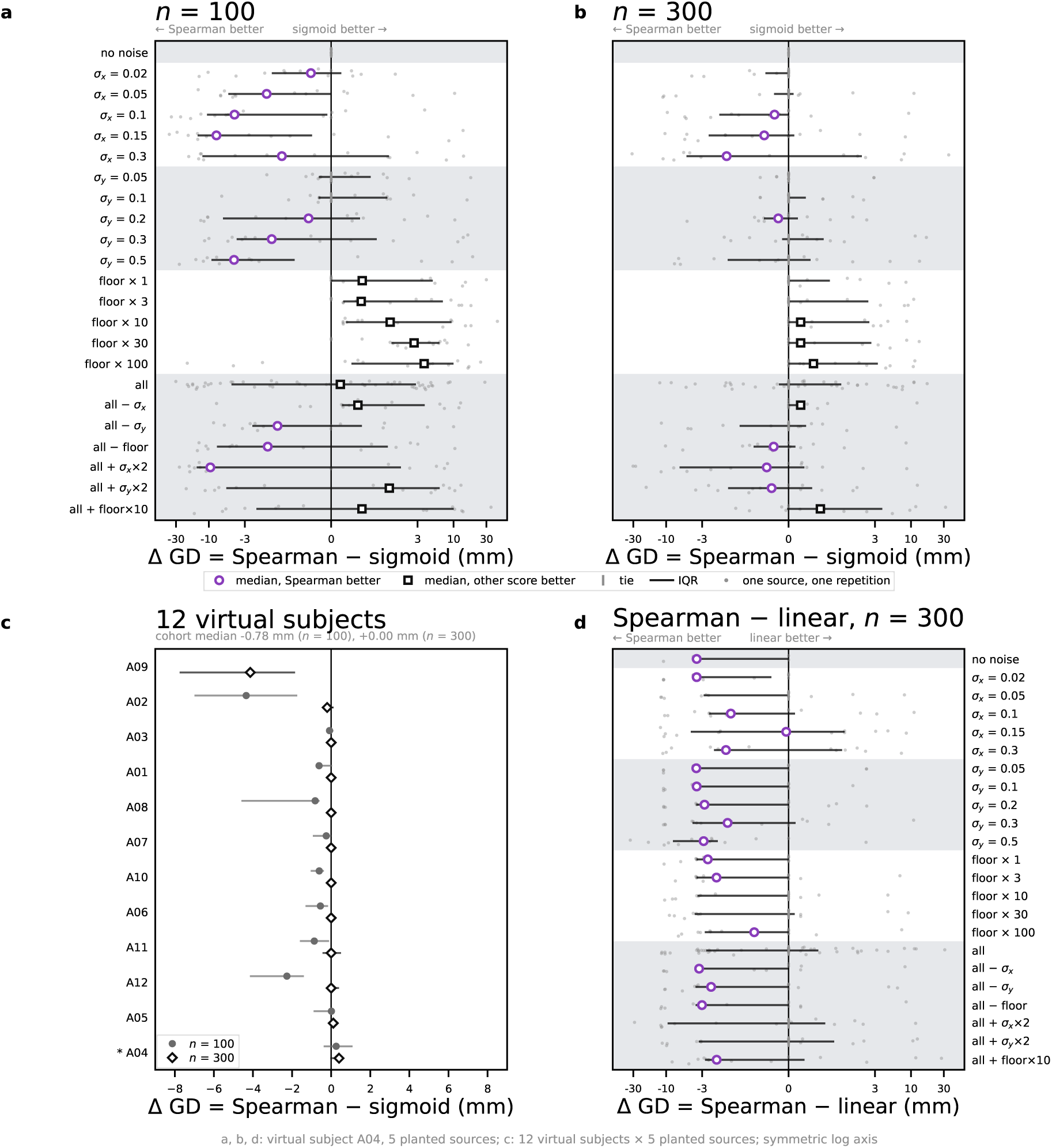
: **Paired difference between the Spearman score and the sigmoid.** Difference in geodesic distance to the planted source, Spearman minus sigmoid; negative values favor Spearman. (a, b) One virtual subject at 100 (a) and 300 pulses (b), one row per noise condition (Appendix D); circle or square, median (circle, Spearman better); bar, interquartile range; gray, individual sources and repetitions. (c) Each of the 12 virtual subjects with all three sources of variability at their calibrated levels. (d) As (b), Spearman minus linear.

**Figure D.3.**
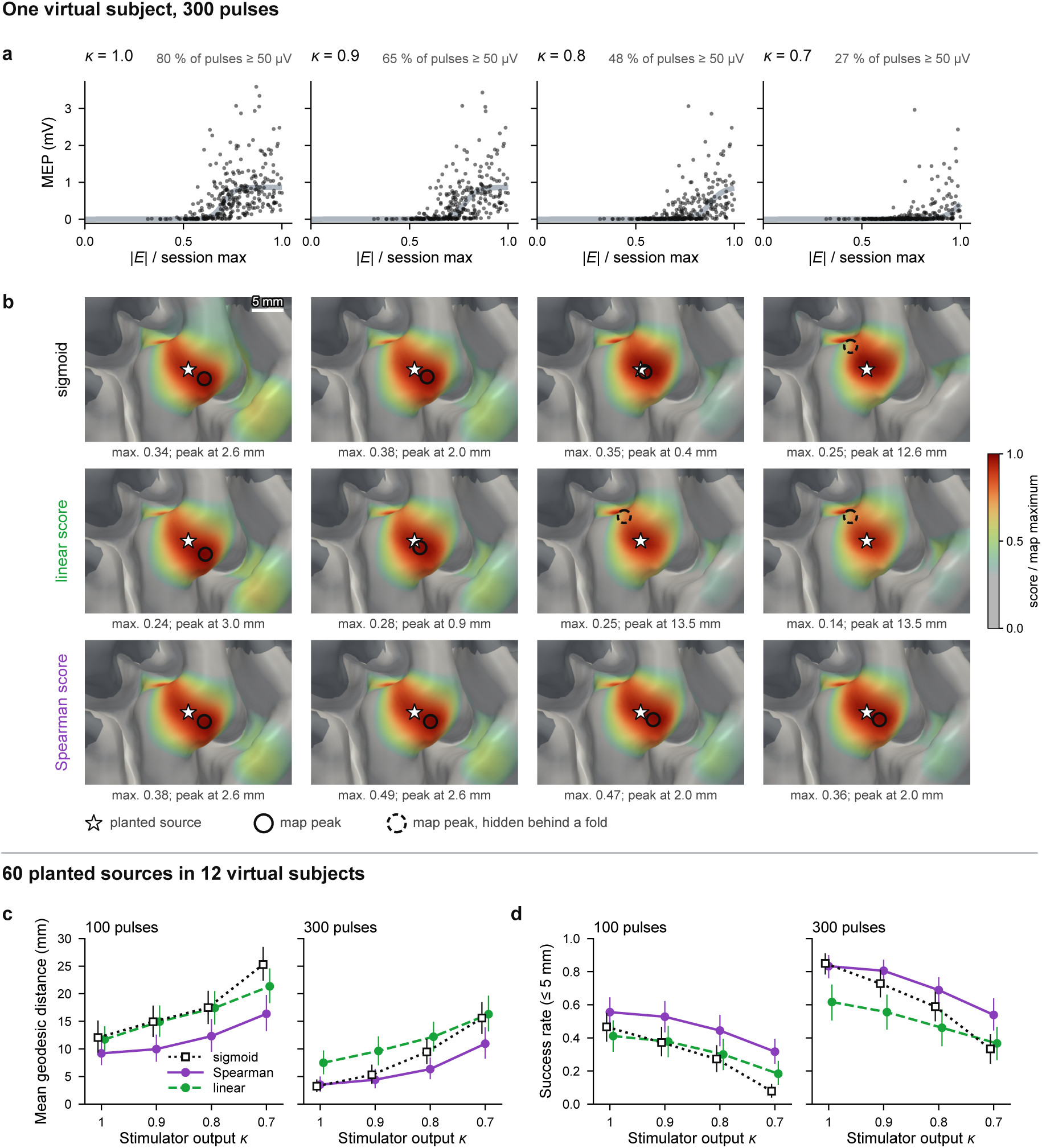
: **Localization at reduced stimulator output.** *κ*, stimulator output relative to the recorded session. (a) MEPs of one virtual subject (300 pulses) against the electric field at the planted source; gray band, the noise-free input–output curve. (b) Sigmoid, linear and Spearman score maps from the same pulses, each scaled to its maximum; below each map, the maximum score and the distance from the map peak to the planted source. (c, d) Mean geodesic distance to the planted source (c) and success rate (peak within 5 mm; d) over 60 planted sources in 12 virtual subjects, at 100 and 300 pulses; bars, 95% confidence intervals.

The three variability sources affect the methods differently (Fig. D.1). Input jitter passes through the steep part of the recruitment curve and reaches the response as heteroscedas-tic noise, which violates the least-squares assumption of the sigmoid fit and degrades it most. The rank score only requires monotonicity and is much less affected. The additive EMG floor degrades the Spearman score only, because the ranks of sub-threshold pulses are then determined by noise, while the baseline parameter of the sigmoid absorbs the floor and its amplitude weighting makes those pulses negligible. The multiplicative gain noise degrades the linear score only, because it acts as heavy-tailed multiplicative noise on the raw amplitudes that the Pearson correlation is computed on.

### D.6 Under-stimulation

We emulated under-stimulation by scaling the drive of every pulse, *u_i_* → *κu_i_*, with *κ* = 1.0, 0.9, 0.8 and 0.7 the stimulator output relative to the recorded session, with the calibrated noise and three paired replays per problem. The share of pulses evoking an MEP of at least 50 µV fell from a median of 0.62 at *κ* = 1 to 0.18 at *κ* = 0.7, and the pulses covered only the lower part of the input–output relation (Fig. D.3a, b).

All three scores lost accuracy as *κ* decreased (Fig. D.3c, d). The Spearman score was the most accurate at every reduced output, and the sigmoid lost the most: at 300 pulses its success rate fell from 0.85 to 0.33, against 0.83 to 0.54 for the Spearman score and 0.62 to 0.37 for the linear score.

